# The self-face captures attention without consciousness: evidence from the N2pc ERP component analysis

**DOI:** 10.1101/2020.01.22.915595

**Authors:** Michał Bola, Marta Paź, Łucja Doradzińska, Anna Nowicka

## Abstract

It is well established that stimuli representing or associated with ourselves, like our own name or an image of our own face, benefit from preferential processing. However, two key questions concerning the self-prioritization mechanism remain to be addressed. First, does it operate in an automatic manner during the early processing, or rather in a more controlled fashion at later processing stages? Second, is it specific to the self-related stimuli, or can it be activated also by other stimuli that are familiar or salient? We conducted a dot-probe experiment to investigate the mechanism behind attentional prioritization of the self-face image and to tackle both questions. The former, by employing a backwards masking procedure to isolate the early and preconscious processing stages. The latter, by investigating whether a face that becomes visually familiar due to repeated presentations is able to capture attention in a similar manner as the self-face. Analysis of the N2pc ERP component revealed that the self-face image automatically captures attention, both when processed consciously and unconsciously. In contrast, the visually familiar face did not attract attention, neither in the conscious, nor in the unconscious condition. We conclude that the self-prioritization mechanism is early and automatic, and is not triggered by a mere visual familiarity. More generally, our results provide further evidence for efficient unconscious processing of faces, and for a dissociation between attention and consciousness.

## Introduction

The ability to recognize oneself is one of the key aspects of self-consciousness. It seems intuitively clear that stimuli representing ourselves, like a self-name or a self-face image, hold a special status in our mind. A classic demonstration of the phenomenon is when attention is automatically drawn to a conversation in which our own name is mentioned - a so-called “cocktail party” effect (Cherry, 1953; Moray, 1959). Multiple experimental studies have subsequently confirmed preferential processing of self-related stimuli (review: Deuve and Bredard, 2011; Sui and Humphreys, 2015; Humphreys & Sui, 2015; Sui and Gu, 2017). Specifically, a range of behavioral paradigms indicate that self-referential stimuli are detected faster and with higher accuracy (Keyes et al., 2010; Sui et al., 2012, 2014; Schafer et al., 2016; Macrae et al., 2018), and are memorized better (Cunningham et al., 2008; Turk et al., 2008; meta-analysis: Symons and Johnson, 1997). EEG studies revealed greater amplitude of the P3 ERP component in response to one’s own name or face as a robust electrophysiological marker of the self-preference effect (Tacikowski and Nowicka, 2010; Ninomiya et al., 1998; Sui et al., 2006; review: Knyazev, 2013). In line, the fMRI studies consistently indicate stronger activations of several brain regions, particularly in the right hemisphere, in response to the self-related stimuli (review: Keenan et al., 2000; Devue and Bredart, 2011; Qin and Northoff, 2011).

Prioritized allocation of attention to the self-related stimuli might be one of the key mechanisms behind the self-preference effect (review: Humphreys and Sui, 2016; Sui and Rotshtein, 2019). Indeed, several studies found that both self-name and self-face automatically capture attention (Tong and Nakayama, 1999; Brédart et al., 2006; Alexopoulos et al., 2012; Yamada et al., 2012; Yang et al., 2013; Alzueta et al., 2020). However, others challenge this conclusion by showing that, first, the effect is not automatic but rather contingent on the task type or availability of attentional resources; and second, that it occurs also for other stimuli that are highly familiar, and thus is not specific to self (Bundesen et al., 1997; Gronau et al., 2003; Harris and Pashler, 2004; Devue and Brédart, 2008; Devue et al., 2009a, 2009b; Keyes and Dlugokencka, 2014; Qian et al., 2015). To further investigate the role of attention in the self-preference effect we have recently conducted a series of studies in which face images - including the subject’s own face - were presented as task-irrelevant distractors in the dot-probe task. Analysis of the N2 posterior-contralateral (N2pc) ERP component - a classic electrophysiological index of covert attention shifts (Eimer and Kiss, 2008; Sawaki and Luck, 2010) - revealed that the self-face automatically and involuntarily attracts attention (Wójcik et al., 2018). Crucially, attention was shifted towards the self-face also when faces were rendered invisible by backward masking and participants were unaware of their identity (Wójcik et al., 2019). Therefore, our findings indicate that identification of the self-face might be performed pre-attentively and pre-consciously.

To elucidate the self-prioritization mechanism it is critical to address whether stimuli representing self are indeed special and processed in a qualitatively different way, or rather the observed effects can be explained by exceptional familiarity with our own name or face. Importantly, in studies focusing on face perception different aspects of familiarity are investigated (review: Ramon and Gobbini, 2018). One aspect is *pre-experimental familiarity*, which assumes that an observer is visually familiar with a given persons’ face and has either extensive knowledge about or an emotional relation with the person. Thus, this aspect is typically investigated using faces of famous people (celebrities) or people close to the participant (e.g. friends or family members). Several studies have shown that such pre-experimentally familiar faces cause attention capture (e.g. Devue and Brédart, 2008; Devue et al., 2009a, 2009b) or ERP effects (Tacikowski et al., 2014) similar to those caused by the self-face. In contrast, *intra-experimental familiarity* is defined as a mere visual familiarity with a given face without extensive knowledge or an emotional component. Previous studies indicate that visually familiar faces might also be processed preferentially and benefit from attentional prioritization, again suggesting an important role of familiarity in the self-preference effect (Tong and Nakaywam, 1999; Tanaka et al., 2006; Tacikowski et al., 2011). However, strong evidence against the familiarity hypothesis is provided by experiments revealing that even abstract stimuli arbitrarily assigned to represent a participant during the experiment benefit from a robust prioritization effect (e.g. Sui et al., 2012, 2014).

Therefore, the present study had three goals. First, we aimed to replicate and extend our previous findings of an automatic attention capture by a consciously and unconsciously processed image of one’s own face (Wójcik et al., 2018, 2019). In line with our previous studies, we used the N2pc component as a primary measure of attention. This was motivated by the fact that ERPs provide a continuous measure of neuronal activity and are able to reveal covert and transient attention shifts. In contrast, the RT index is an aggregate measure of a whole chain of processes between perception and a manual response, and thus might not be sensitive enough to indicate transient attention shifts (see: Kappenman et al., 2014). Our second aim was to clarify the role of visual familiarity in the observed effect by testing whether a visually familiar face will attract attention in a similar manner as the self-face. Participants were therefore familiarized with a previously unfamiliar face image, which was then repeatedly presented in each trial of the “familiar” condition (similarly to the self-face in the “self” condition). Thus, it is important to emphasize that in the present manuscript the term “familiarity” refers to the *intra-experimental (visual) familiarity.* Our third aim was to investigate whether attentional prioritization of both self- and familiar-face would change across the time-course of the experiment. We hypothesized that the familiar face will capture attention more strongly in the second phase of the experiment, as it’s familiarity will increase across trials (in line with: Tong and Nakaywam, 1999; Tanaka et al., 2006; Tacikowski et al., 2011). However, we did not expect such an effect for the self-face, which exhibits a ceiling-level familiarity and thus further experimental presentations are unlikely to increase its effect (Tong and Nakaywam, 1999; Tacikowski et al., 2011).

## Methods

### Participants

We collected data of 29 participants (16 females, *M* = 23 years, *SD* = 3 years, range: 20-29 years, 1 lefthanded). They all declared normal or corrected-to-normal vision and no history of psychiatric or neurological disorders.

Data of 6 additional participants were collected but excluded from the analysis: EEG data of 1 subject has not been properly saved, electro-oculographic (EOG) signal of 2 participants was not properly recorded and could not be used in the analysis, and 3 participants were excluded due to an insufficient number of epochs remaining after the EEG preprocessing procedure (detailed criteria are described in the “EEG recording and analysis” section).

All experimental procedures were approved by the local Research Ethics Committee at the SWPS University, Warsaw, Poland. All participants provided written informed consent and received monetary compensation for their time (100 PLN = c.a. 25 EUR).

### Stimuli

A group of 35 participants were initially recruited for the project (20 females, 15 males). All participants were first invited to the lab to have a photograph of their face taken in a standardized environment (the same background and lightning conditions) with a 12-megapixel digital camera (Canon Powershot SX130). Participants were asked to maintain a neutral facial expression when photographed. Next all photographs were cropped to include only the face oval (i.e. without hair) and then normalized in terms of luminance, contrast, and grey-scaled colour using Photoshop CC 2018 (Adobe, San Jose, CA). Further, 35 “masks” were created in order to use them to backward mask the face images during the experiment. Masks were created by randomly relocating the key elements of each face (eyes, nose, mouth) in order to disturb the classic feature configuration of faces and prevent recognition (similarly to our previous work: Wójcik et al., 2019).

Next participants were invited for an EEG experimental session. For each subject, the following stimuli were used: an image of his/her own face; one face chosen from the pool of same gender faces to constitute an intera-experimentally familiar face; and 10 faces chosen randomly from the pool of same gender faces, which were used as control stimuli (other faces). Masks used for backward masking were also gender-matched for each participant (i.e. male participants were presented only with male faces, which were masked with masks created from male faces only).

### Procedure

The experimental procedure was written in the Presentation software (Neurobehavioral Systems, Albany, CA, USA) and presented on a FlexScan EV-2450 (Hakusan, Ishikawa, Japan) screen through an Intel Core i3 computer. Participants were sat comfortably in a dimly lit room with a viewing distance of 70 cm, which was maintained by a chinrest. The experimental procedure started with a display providing participants with information about the structure of a trial, the instruction to focus on detecting and manually responding to a target dot, and information that displayed faces are distractors and thus should be ignored. Next, participants were presented with the face image chosen to constitute an intra-experimentally familiar face. The image was accompanied with a short fictional story, with which we aimed to establish a basic representation of the person: “This is Ania/Tomek. She/He’s from Warsaw. She/He is 23 and studies economy. She/He works at a cafe in the Wola district and likes visiting national parks”. Participants were asked to familiarize themselves with the face and the provided information, and press a “continue” button whenever they feel ready. In the next screen both the self-face and the intra-experimentally familiar face were presented together, with an information that different faces will be presented during an experiment, including participants own face and “Ania’s/Tomek’s” face, but the subject’s task is to focus on the target dot and ignore appearing faces. With this additional presentation we aimed to further familiarize participants with features of both self-face and familiar-face images, so that the familiar face would not exhibit an advantage.

A block design was used in the experiment. The tasks were always presented in the following fixed sequence: masked dot-probe task, masked identification task, unmasked dot-probe task, unmasked identification task. Masked (unconscious) condition always proceeded the unmasked (conscious) condition as we did not want conscious presentations to increase participants’ familiarity with the physical features and consequently lower the perception threshold in the masked trials (e.g. Lamy et al., 2017). Within each block there was a self-face block and a familiar-face block (200 trials each, with a break after 100 trials provided for participants comfort). It was randomly chosen whether the self- or familiar-face block would be presented first.

#### Dot-probe task

In the self-face blocks of the dot-probe task (and analogously in the familiar-face blocks) the self-face (familiar-face) image was presented in each of 200 trials and was always paired with one of 10 “other” faces (thus, each of these faces appeared 20 times).

All stimuli were presented against black background. A dot-probe trial started with a presentation of a fixation cross (subtending 0.3° x 0.3° of the visual angle) at the centre of the screen. The fixation cross remained onscreen for the duration of the trial. After either 750 ms or 1250 ms (the jitter was chosen randomly) a pair of faces were presented bilaterally for 32 ms. Faces subtended 6.2° × 7.9° of visual angle, with their inner edge 3° left and right from the fixation cross. In the masked blocks the faces were followed by backward masks, which remained on a screen for 50 ms. In each trial two masks were chosen randomly from the pool of all gender-matched masks. In the unmasked blocks a blank screen was displayed for 50 ms. Next, a target asterisk subtending 0.3° x 0.3° of the visual angle was presented for 150 ms in the location of the centre of either the self-face/familiar-face (congruent trial) or the other face (incongruent trial). Within each block half trials were congruent and half were incongruent, and their order was random. Participants were asked to indicate - by pressing one of two buttons using index fingers of their left or right hand - the dots’ presentation side (left or right). Participants were asked to respond as quickly and accurately as possible. The response time was limited to 3000 ms and the next trial started immediately after the manual response.

#### Identification task

In the self-face blocks of the identification task (and analogously in the familiar-face blocks) the self-face (familiar-face) image was presented in half of the 200 trials and always paired with one of 10 “other” faces. In the other half of the trials, two “other” faces were displayed. The structure of the trial was the same as in the dot-probe task, but the target dot was not presented. Instead, participants were asked to make a forced-choice whether their own (in the self-face identification task) or the familiar-face (in the familiar-face identification task) was presented on a given trial. Participants were informed that the response time is unlimited and asked to respond as accurately as possible by pressing one of two buttons of the response pad.

### Analysis of behavioral data

All analyses of behavioral data were conducted using custom-made Python scripts. Analysis of the dot-probe task data tested whether reaction times (RT) of manual responses to the target dot differ between two types of trials: those in which dots followed the potentially attention-grabbing stimulus (self- or familiar-face), and those in which dots followed the neutral stimulus (other face). Included in the analysis were only responses that were correct and occurred between 200 ms and 1600 ms after dot presentation. For each subject and condition a median RT was calculated and used in the statistical analysis. Accuracy of responses to the target dot was calculated as a percentage of correct responses. The obtained values are presented in the Results section, but due to ceiling level performance in the majority of participants this measure was not analysed statistically.

Sensitivity index *(d’)* was calculated from the identification task data to evaluate whether participants were able to distinguish between the target (presence of the self- or familiar-face) and the “noise” stimuli (absence of the self- or familiar-face) in the identification task. Hits and false alarms equal to 0 or 1 for each subject were replaced using the log-linear rule, the least biased method of correcting extreme values (Stanislaw and Todorov, 1999).

### EEG recording and analysis

EEG signal was recorded with 64 Ag-AgCl electrically shielded electrodes mounted on an elastic cap (ActiCAP, Munich, Germany) and positioned according to the extended 10-20 system. Vertical (VEOG) and horizontal (HEOG) electrooculograms were recorded using bipolar electrodes placed at the supra- and sub-orbit of the right eye and at the external canthi. Electrode impedances were kept below 10 kΩ. The data were amplified using a 128-channel amplifier (QuickAmp, Brain Products, Enschede, Netherlands) and digitized with BrainVisionRecorder® software (Brain Products, Munich, Germany) at a 500 Hz sampling rate. The EEG signal was recorded against an average of all channels calculated by the amplifier hardware.

EEG and EOG data were analyzed using EEGlab 14 functions and Matlab 2016b. First, all signals were filtered using high-pass (0.5 Hz) and low-pass (45 Hz) Butterworth IIR filter (filter order = 2; Matlab functions: *butter* and *filtfilt).* Then data were re-referenced to the average of signals recorded from left and right earlobes, and down-sampled to 250 Hz. All data were divided into 800 epochs (200 epochs per dot-probe condition; conditions: familiar masked, self masked, familiar unmasked, self unmasked). Epochs were created with respect to the onset of face-images ([-200, 800] ms) and baseline-corrected by subtracting the mean of the pre-stimulus period ([-200, 0] ms). Further, epochs were rejected based on the following criteria: i) when reaction-time (RT) of the manual response to the target dot was < 100 ms or > 800 ms (M = 11.5±2.7; range: [0, 46] epochs per subject); ii) when activity of the HEOG electrode in the time-window [-200, 500] ms exceeded −40 or 40 uV (M = 118.2±21.3; range: [11, 391] epochs); iii) when activity of the P7 or P8 electrode in the time-window [-200, 800] ms exceeded −60 or 60 uV (M = 5.7±2.9; range: [0, 80] epochs). Thus, after applying the described criteria the average number of analyzed epochs per subject was: 664.4±24, range: [372, 788]. A subject was excluded if the number of epochs in any of four conditions was < 50. This criterion resulted in excluding 3 participants out of 32 (but additional 3 participants were excluded due to other criteria, as described in the “Participants” section). The numbers of epochs provided above were calculated based on the final sample of 29 participants. We found moderate or anecdotal evidence indicating no difference between conditions when compared in terms of number of retained epochs (Bayes Factor < 0.65 for all comparisons).

Next, each EEG-EOG data-set was decomposed into 50 components using Independent Component Analysis as implemented in the EEGlab *pop_runica* function. To remove residual oculographic artefacts from the data the following procedure was used: the time-course of each component was correlated with time-courses of HEOG and VEOG electrodes and a component was subtracted from the data if either value of the Spearman correlation coefficient exceeded −0.3 or 0.3. Using this procedure M = 3.2±0.3; range: [2, 6] components per subject were removed.

After applying the described preprocessing steps, data were divided with respect to the condition and presentation side of the self/familiar face. To calculate the N2pc component we used the P8 and P7 electrodes. Specifically, when the self/familiar face was presented on the left side, P8 was the contralateral electrode and P7 was the ipsilateral electrode. When the self/familiar face was presented on the right side, P7 was the contralateral electrode and P8 was the ipsilateral electrode. For each condition contralateral and ipsilateral signals were first concatenated and then averaged, to obtain waveforms presented in **Fig. 1.** To obtain estimates of the early and late parts of the N2pc component for the statistical analysis the waveforms were averaged across time within, respectively, 200 - 300 ms and 300 - 400 ms time-windows. The timewindows are consistent with our previous study (Wójcik et al., 2019).

**Figure 1.**
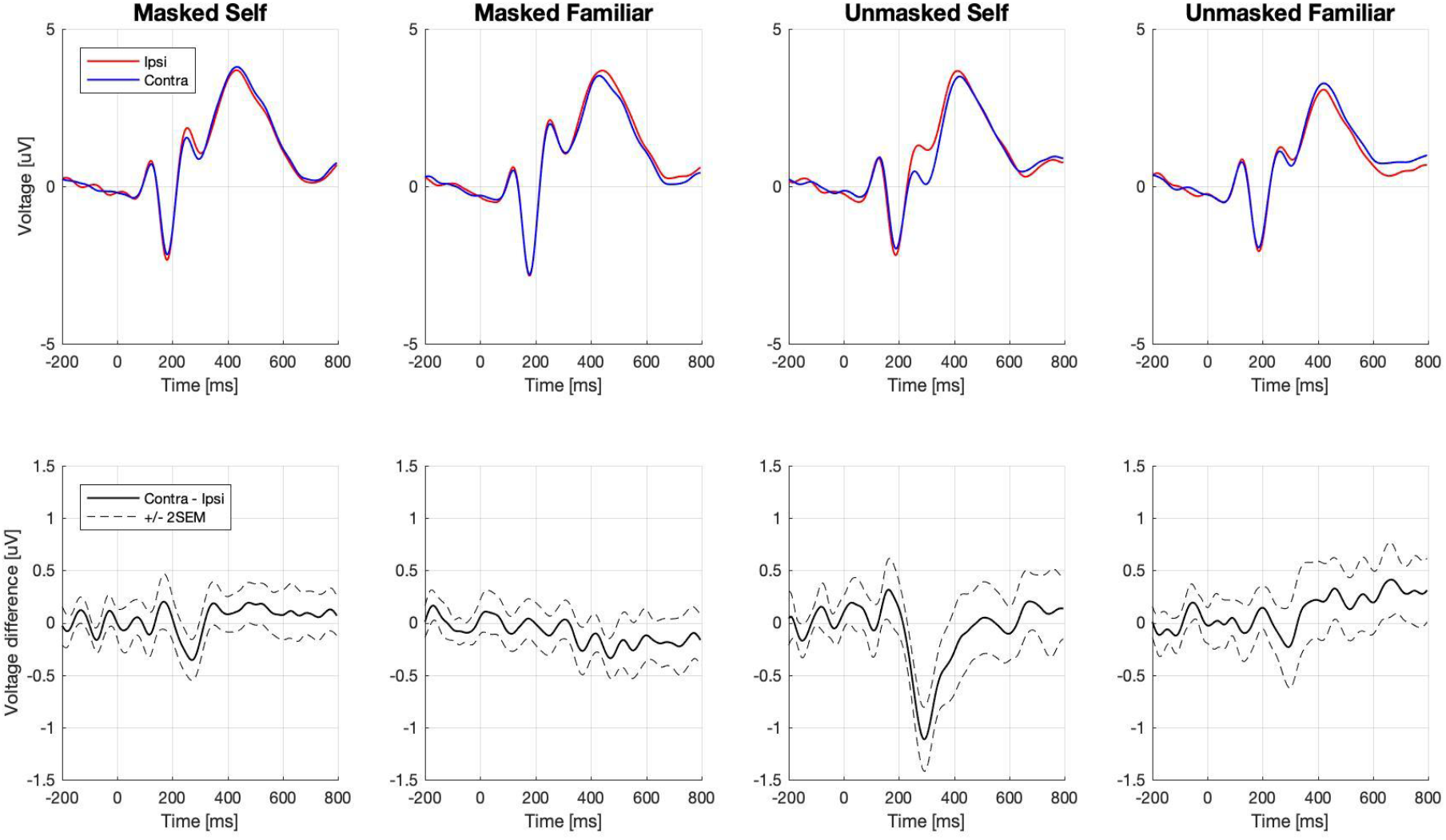
Event related potentials in the dot-probe task. Electrodes P7/P8 were chosen for the analysis. Waveforms from electrodes ipsi- and contra-lateral to the self- and familiar-face images are presented in the top row. Difference waveforms (i.e. contra - ipsi-lateral) are presented in the bottom-row.

### Statistical analysis

All statistical analyses were conducted in the JASP software and cross-checked with Statcheck (http://statcheck.io/index.php). The values are reported as Mean (*M*±SEM) with 95% confidence interval (CI), unless stated otherwise. Because the dot-probe accuracy values exhibited highly right-skewed distributions they are reported as Median (Mdn±quartile deviation).

RT and N2pc effects were investigated with repeated measures ANOVA models. Masked and unmasked conditions were analysed in separate models, as the order of their presentation was fixed (with masked condition always preceding the unmasked condition). Specific to the analysis of RT was the factor of *congruency*, defined by the dot presentation side with respect to the self/familiar face (congruent vs. incongruent trials). Specific to the electrophysiological analysis were two factors: *N2pc*, defined the side on which ERP activity was recorded, with respect to the self/familiar face (ipsi- vs. contra-lateral); and the *time-window* in which it was recorded (early vs. late N2pc). The factor of a *stimulus* type (self face vs. familiar face) was included in all three models. Further, within each condition all trials were divided into first and second half (100 trials each) and this factor *(half)* was also included in four three models in order to test the hypothesis that the familiar-face captures attention only after reaching a certain level of familiarity. ANOVA results were reported in tables or as *F*(df) and the partial-eta squared (an indicator of the effect size) was reported as 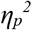. In case of a significant interaction we investigated simple effects to provide further insight into the data structure.

The *d’* index, estimated based on the identification task data, was at the group level tested against 0 and compared between conditions. Data distribution was first tested with the Shapiro-Wilk test. Two-tailed one-sample and paired-sample t-tests were used and reported as *t*(df), while the effect-size established with Cohen’s d was reported as *d*. Correlation between *d’* and N2pc amplitude (calculated as difference between contra- and ipsi-lateral activity) was calculated using one-tailed Pearson’s correlation test (*r*). In case of *d’* analyses the traditional statistical approach was complemented with Bayesian statistics, which allow testing for the lack of differences between variables. We have computed the Bayes Factor (BF10) which is defined as the ratio of the probability of observing the data given the alternative hypothesis (H1) is true to the probability of observing the data given the null hypothesis (H0) is true. We used Bayesian one-sample and paired t-tests, all with medium prior scale (Cauchy scale 0.707), in order to test for differences and lack of differences between means. We also performed Bayesian Pearson’s correlation to relate the *d’* values and the ERP amplitudes. In the results section we provide interpretations of the BF according to Wagenmaker et al. (2018).

In all conducted statistical analyses *p* value < 0.05 was considered to indicate a statistically significant effect. Because 4 ANOVA models were conducted in this study (2 models with 3 factors, and 2 models with 4 factors) only significant main effects and interactions are reported in the Results section. Details of all main effects and interactions are presented in Tables included as Supplementary Materials.

### Data availability statement

All data (including raw EEG data) and scripts used for presentation of stimuli and for analysis will be shared by authors per request.

## Results

### Identification task

To evaluate participants’ ability to recognize the self-face and the familiar face the *d’* sensitivity index was calculated. In the unmasked condition *d’* values were high for both the self-face *(M* = 2.79±0.206; *t*(28) = 13.56, *p* < 0.001; BF > 1000) and the familiar face (*M* = 2.61±0.193; *t*(28) = 13.56, *p* < 0.001; BF > 1000), indicating that despite short display times participants perception was fully conscious. In the masked condition relatively low mean *d’* values were observed, which indicate that perception was highly degraded, however, comparisons to 0 indicate that performance was above chance-level in both self-face (*M* = 0.31±0.073; *t*(28) = 4.22,*p* < 0.001; BF = 123) and familiar-face conditions (*M* = 0.18±0.055; *t*(28) = 3.42, *p* = 0.002; BF = 18). Further, comparing *d’* between the self- and familiar-face we found no difference between conditions (*t*(28) = 1.31, *p* = 0.20; BF = 0.42, null hypothesis 2.3 more likely).

### Dot-probe task - behavioral results

High accuracy values were observed in all dot-probe task conditions. For congruent and incongruent trials, respectively, the values were as follows: masked self-face (*Mdn* = 96±3% and *Mdn* = 95±3.5%); masked familiar-face (*Mdn* = 96±2.5% and *Mdn* = 97±2.5%); unmasked self-face (*Mdn* = 95±2% and *Mdn* = 95±2.5%); and unmasked familiar-face (*Mdn* = 96±3% and *Mdn* = 96±3%). Statistical analysis of the accuracy values was not performed because the majority of participants exhibited ceiling level performance.

Therefore, reaction times (RT) of manual responses to target dots were analyzed as a behavioral index of attention shifts in the dot-probe task. Specifically, we investigated whether RT was shorter when the target dot followed a potentially attention-grabbing self- or familiar-face image (i.e. congruent trials), in comparison to trials when it followed a control stimulus (i.e. incongruent trials). However, the estimated values suggested no difference between congruent and incongruent trials, neither in the masked (self-face: *M* = 368.92±10.74 vs. *M* = 368.42±10.26; familiar-face: *M* = 366.96±8.41 vs. *M* = 369.42±8.66), nor in the unmasked condition (self-face: *M* = 337.45±6.79 vs. *M* = 341.60±7.82; familiar-face: *M* = 338.11±7.07 vs. *M* = 340.98±8.57). Indeed, the three-way repeated-measures ANOVA indicates no significant main effects or interactions, neither in the masked, nor in the unmasked condition (**Tables S1, S2**).

### Dot-probe task - N2pc results

Electrophysiological data were collected during the dot-probe task to analyze the N2pc component, which is a classic index of covert attention shifts (Eimer and Kiss, 2008; Sawaki and Luck, 2010). N2pc is considered to occur when an ERP recorded contra-laterally to the stimulus exhibits lower amplitude than an ERP recorded ipsi-laterally (**Fig. 1**). Consistently with our previous studies, we analyzed the early (200-300 ms) and late (300-400 ms) parts of the N2pc component (Wójcik et al., 2019). Further, in the present study we also investigated whether the N2pc component is affected by the number of stimulus presentations (by comparing the first and second half of trials within each block).

In the masked conditions the four-way repeated-measures ANOVA (**Tables S3, S4**) revealed a significant interaction of N2pc, stimulus type, and ERP time-window (*F*(1, 28) = 7.06, *p* = 0.013, 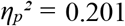). Analysis of simple effects indicates that the self-face stimulus evoked the early N2pc (ipsilateral amplitude: *M* = 0.87±0.45 μV; contralateral: *M* = 1.09±0.41 μV; *F*(1, 28) = 7.06, *p* = 0.013, 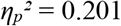). There was no evidence that the late N2pc was evoked by the self-face (ipsilateral: *M* = 2.17±0.48 μV; contralateral: *M* = 2.10±0.48 μV; *F*(1, 28) = 0.66, *p* = 0.424, 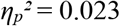). Similarly, no N2pc was observed in response to the familiar-face, neither in the early (ipsilateral: *M* = 1.11±0.52 μV; contralateral: *M* = 1.24±0.48 μV; *F*(1, 28) = 2.27, *p* = 0.143, 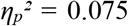), nor in the late time-window (ipsilateral: *M* = 1.88±0.48 μV; contralateral *M* = 2.30±0.47 μV; *F*(1, 28) = 2.81, *p* = 0.105, 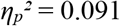).

In the conscious (unmasked) condition the conducted ANOVA model (**Tables S5, S6**) revealed a significant interaction between N2pc and stimulus type (*F*(1, 28) = 14.15, *p* <0.001, 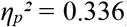), but the time-window had no impact on this effect. Analysis of simple effects shows that N2pc component was evoked by the self-face (ipsilateral: *M* = 0.76±0.41 μV; contralateral: *M* = 1.33±0.39 μV; *F*(1, 28) = 24.90, *p* < 0.001, 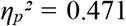), but there was no evidence it was evoked by the familiar-face (ipsilateral: *M* = 1.10±0.49 μV; contralateral: *M* = 1.03±0.52 μV; *F*(1, 28) = 0.248, *p* = 0.622, 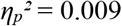). Of note, we did not find any difference between the first and second half of the trials, neither in the masked, nor in the unmasked condition.

### Correlation of perceptual sensitivity and N2pc

To evaluate whether the above chance performance in the masked identification task might have driven the unconscious attention capture effect we correlated *d*’ and early N2pc. For the masked self-face we found anecdotal evidence for lack of correlation when *d*’ was correlated with early N2pc (*r* = −0.22; *p* = 0.24; BF = 0.44). However, a significant correlation between *d*’ and early N2pc was found for the unmasked self-face presentations (*r* = −0.44; *p* = 0.019; BF = 3.15), which closely replicates our previous finding (Wójcik et al., 2019) and suggests that in the unmasked condition the ability to recognize one’s own face better was related to stronger attention capture.

## Discussion

The ability to recognize oneself is fundamental for creating and maintaining a coherent personal identity (Sui and Gu, 2017). Preferential allocation of attention to stimuli representing or associated with ourselves might be a key mechanism facilitating self-recognition (Sui and Rotshtein, 2019). In the present study we analyzed the N2pc ERP component - a classic marker of covert attention shifts (Eimer and Kiss, 2008) - and found that the self-face image indeed captures attention automatically and involuntarily. Most importantly, the self-face attracted attention even when processed outside of awareness. Our second aim was to investigate whether a mere visual familiarity with our own face might explain the observed self-preference effect. However, we did not find any evidence that attention was captured by a face that was visually familiar and repeatedly presented to participants, neither in the conscious, nor in the unconscious condition. Below we discuss our findings in the context of previous research on the self-preference effect, the role of familiarity, and the relation between attention and perceptual consciousness.

### The role of attention in perception of self-related stimuli

A significant body of evidence indicates that self-related information benefits from preferential processing at the late, cognitive processing stages. This is suggested for instance, by greater amplitude of the P3b ERP component evoked by the self-name or face (e.g. Tacikowski and Nowicka, 2010; Doradzińska et al., 2020; review: Knyazev, 2013) or better memory of self-related stimuli (meta-analysis: Symons and Johnson, 1997). However, it is controversial whether the self-prioritization effect occurs already at the perceptual processing stages and in an automatic manner. Automatic effects can be defined as unintentional, unconscious, and not dependent on the availability of perceptual or cognitive resources. While several studies suggest that the self-prioritization meets these criteria (Tong and Nakayama, 1999; Brédart et al., 2006; Alexopoulos et al., 2012; Yamada et al., 2012; Yang et al., 2013; Doradzińska et al., 2020), others claim the observed effects do not comply with this definition of automaticity (Bundesen et al., 1997; Gronau et al., 2003; Harris and Pashler, 2004; Devue and Brédart, 2008; Devue et al., 2009a, 2009b; Keyes and Dlugokencka, 2014; Qian et al., 2015). Here we support the former view by showing that the self-face attracts attention even when displayed as a task-irrelevant distractor and when its perception is subliminal, which suggests the effect is unintentional and preconscious. What remains to be investigated is the role of perceptual resources in self-face processing, which has not been explored here and thus will be a goal of future studies.

Controversies regarding the self-preference effect might be potentially explained by variability in types of self-related stimuli used across experiments. While the “cocktail party” effect is considered a classic demonstration of the early and preattentive ability to identify an auditorily presented self-name (Cherry, 1953; Moray, 1959; but see the criticism of Lachter et al., 2004), the visually presented self-name is typically related with relatively late brain activations, specifically the P3b component (Doradzińska et al., 2020; review: Knyazev, 2013). In contrast, the visual image of the self-face seems to also cause earlier effects, likely reflecting perceptual or attentional stages of processing. Several studies found ERP signatures of the self-face processing around 250 ms (Caharel et al. 2002; Sui et. al 2006; Tanaka et al., 2006; Alzueta et al., 2019), or even around 170-200 ms after the stimulus onset (e.g. enhanced N170 component; Keyes et al., 2010). These observations are in line with our data, suggesting that the self-face is identified and prioritized already around 200 ms after the stimulus onset.

In dot-probe studies two measures - the RT effect (i.e. shorter RT in congruent, in comparison to incongruent trials) and the N2pc effect (i.e. more negative N2 component contralateral to the stimulus, in comparison to the ipsilateral N2) - are considered equivalent indexes of attention shifts. Importantly, here we found a dissociation between these two measures, i.e. presence of the N2pc effect, but lack of the RT effect in the self-face condition. This finding is in line with several previous dot-probe studies using the self-face (Wójcik et al., 2018, 2019) or emotional images as stimuli (Kappenman et al., 2014, 2015; Furtak et al., 2020). What can account for the observed dissociation is that the RT of a manual response reflects an aggregate outcome of a whole chain of processes occurring between a stimulus presentation and a response; and thus the RT index might not be sensitive enough to indicate transient and automatic attention shifts. Further, considering that detection of the probe is typically very simple, it might not be impaired even in the presence of an attention grabbing distractor. In contrast, ERPs provide a continuous measure of neuronal engagement and thus can reveal covert attention shifts not manifested in overt behavior. Indeed, Kappenman and colleagues (2014) demonstrated that the RT index exhibits poor internal reliability in the dot-probe task (and consequently its external validity must be also poor), while reliability of the N2pc is moderate. Thus, an important goal for future studies will be defining the functional role of the N2pc component and its relation to behavior (e.g. Zivony et al., 2018).

### Unconscious processing of the self-face

The present study demonstrates that the self-face captures attention, even when processed outside of awareness, and thus replicates our recently published finding (Wójcik et al., 2019). Our results are in line with several previous observations, which collectively suggest an automatic and preattentive nature of the self-preference effect. First, the subliminally presented self-name can cause both priming (Pannese and Hirsch, 2010; Pfister et al., 2012) and interference effects (Alexopoulos et al., 2012). Second, self-related stimuli, both names and faces, are boosted into awareness in the attentional blink (AB) or continuous flash suppression (CFS) paradigms (Shapiro et al., 1997; Geng et al., 2012; Macrae et al., 2017; but see: Stein et al., 2016). Third, Sui and colleagues (2012) found that low-intensity stimuli are detected more accurately when related to self, which also suggests an effect at the early and presumably preconscious processing stage (but see: Tacikowski and Ehrsson, 2016). Finally, the self-name modulates amplitude of the late P3b ERP component even when presented subliminally (Doradzińska et al., 2020). All these studies support the notion that self-related information can be identified and prioritized without perceptual awareness.

Of note, our analysis of the dot-probe task data (N2pc component) indicates that the self-face was attentionally prioritized, but analysis of the identification task data (*d’* index) suggests that the self-face did not gain preferential access into awareness (in comparison to the familiar-face). While these two findings might be considered contradictory, we argue they provide support for the putative dissociation of attention and perceptual awareness. Further, that the self-face was not boosted into awareness might be also considered at odds with the above mentioned AB or CFS studies. However, in AB and CFS paradigms lack of awareness is mainly caused by lack of attentional resources (i.e. preconscious processing, according to Kouider and Dehaene, 2007) and therefore self-related stimuli, with their ability to attract attention, robustly break into awareness. In contrast, in the backward-masking paradigm lack of awareness is caused by weakening the bottom-up input and/or disturbing the top-down feedback processing (i.e. subliminal processing; Kouider & Dehaene, 2007), and thus even stimuli that are attentionally prioritized are unlikely to gain access to awareness. Of note, s similar result has been found in our recent study, in which a backward-masked self-name increased the amplitude of the P3b ERP component (in comparison to an unfamiliar name) but did not gain preferential access to awareness (Doradzińska et al., 2020).

An emerging consensus concerning the relation between attention and consciousness suggests a partial dissociation of the two mechanisms. Specifically, at least minimal attention is considered necessary for perceptual consciousness, but consciousness seems not necessary for attentional selection mechanisms to work (Van Boxtel et al., 2010; Cohen et al., 2012; Pitts et al., 2018). Indeed, a growing body of evidence suggests that stimuli processed outside of awareness might nevertheless capture attention (Woodman and Luck, 2003; Ansorage and Heumann, 2006; Hsieh et al., 2011; Lamy et al., 2015). However, previous experiments typically used simple singleton stimuli, while our work demonstrates images of faces - relatively complex and naturalistic stimuli - can be also prioritized without awareness. Importantly, while there is evidence that emotional expression of a face can be detected unconsciously (review: Axelrod et al., 2015; Hedger et al., 2016), it is still being debated whether an identity of a face can be processed without awareness. The results reported so far indicate that identity of an pre-experimentally familiar face (of a famous or personally close person) can be processed unconsciously (Henson et al., 2008; Kouider et al., 2009; Gobbini et al., 2013), but recognizing identity of an unfamiliar face requires awareness (Moradi et al., 2005; Stein and Sterzer, 2011; Axelrod and Rees, 2014). This suggests that unconscious processing is based on activation of a pre-established representation, and our results from both self-face and familiar-face conditions, can be interpreted as supporting this conclusion.

### The role of visual familiarity

Familiarity can be understood as a continuum, with several aspects - like presence and the strength of the emotional component, the amount of information we have, or just the degree of mere visual familiarity - contributing to the general level of familiarity (review: Ramon and Gobbini, 2018). In the present study we focused on the role of *visual (intra-experimental) familiarity*, which was established by presenting an initially unfamiliar face before the experiment and then repeated presentation during the experiment. Thus, the intra-experimental familiarity constitutes the “weakest” form of familiarity, but it allows the most precise experimental control and sets a lower-bound on the effects that other forms of familiarity might cause.

Here we found that a visually familiar face did not capture attention, neither in conscious, nor in unconscious condition. This is important for interpreting data from the self-face condition, as it indicates that mere visual familiarity with our own face is not the main factor causing attentional prioritization. However, our finding seems to disagree with Tong and Nakayama (1999), who revealed that reaction times in a search task decreases rapidly across trials when the unfamiliar face constitutes a target, but there is no improvement in case of the self-face (for which performance exhibits a ceiling level from the beginning). In line, Tanaka et al. (2006) asked participants to detect a previously unfamiliar face and found that amplitude of the N250 - an ERP component indexing familiarity - increased across trials (but again, processing of the self-face did change over time). However, in both studies the “familiar” face was defined as a to-be-detected target, and thus it is likely that participants had to maintain this face in working-memory to perform the task (which, by itself, might cause an automatic attention capture; see: Downing, 2000). In contrast, in our study faces were presented not as targets, but as task-irrelevant distractors which participants were supposed to ignore. This might thus be the main reason why we did not observe any “learning effect” of a familiar face.

### Limitations and conclusions

A possible limitation of our study is that even though the *d’* values observed in the masked conditions were low (self-face: M = 0.31±0.073; familiar-face: M = 0.18±0.055), they were statistically greater than 0 and thus indicate above chance-level performance. One can therefore claim that perception was highly degraded but not fully unconscious, and that the “unconscious” attention capture was caused by the residual awareness. We present three arguments to mitigate this concern. First, N2pc amplitude was not correlated with d’, neither in the masked self-face, nor in the masked familiar-face condition. Second, d’ was estimated based on the identification task data, in which self- and familiar-face were considered task relevant targets. However, in the dot-probe task faces were presented as task-irrelevant distractors, and thus participants were likely less perceptually sensitive to faces in the dot-probe than in the identification task. Third, it is not known to what extent the above chance-level *d’* might indicate an unconscious “blind-sight” type of processing, which has been demonstrated in case of face perception (review: Axelrod et al., 2015).

In conclusion, we found that the self-face robustly captures attention when presented consciously and unconsciously. This finding adds to the body of evidence suggesting an early and automatic prioritization mechanisms for self-related information. However, we did not find any evidence that the visually familiar face was prioritized. In general, our study demonstrates that complex stimuli might be processed and capture attention without consciousness, providing further evidence that attention and consciousness can be dissociated.

## Conflict of interest

The authors declare no competing interests.

## Acknowledgements

This study was funded by the National Science Center Poland grants (2018/29/B/HS6/02152 to MB, and 2018/29/B/HS6/00461 to AN).

## Contributions

Conceived study: MB, AN; Designed study: MB, MP, ŁD, AN; Collected data: MP; Analyzed data: MB, MP, ŁD; Drafted manuscript: MB; Revised manuscript: MB MP, ŁD, AN.

## Notes

### Competing Interest Statement

The authors have declared no competing interest.

### Summary of Updates

The most important improvement is that we used ANOVA models to analyze our data (instead of simple t-tests), and all our conclusions are now based on results of these models. Our main conclusion regarding attention capture by the consciously and unconsciously processed self-face has not changed. However, we did not find any convincing evidence in favour of attention capture by the visually familiar face, which is in line with suggestions of the Reviewers that the evidence found in the previous version of the analysis was relatively weak and perhaps incidental. Therefore, the discussion section was revised accordingly.

