## Supplementary Materials for "The self-face captures attention without consciousness: evidence from the N2pc ERP component analysis"

**Table S1.** Three way repeated measures ANOVA on RT data collected during the masked dot-probe task. The 2x2x2 design involves: *congruency* (the target presentation side congruent vs incongruent with respect to the self-/familiar-face presentation side), *stimulus* (self-face vs familiar-face), and *half* (first vs second half of trials within a block).

| source | Sum of Squares | df | Mean Square | F | p | $\eta_p^2$ |
|---|---|---|---|---|---|---|
| congruency | 209.000 | 1 | 209.000 | 1.787 | 0.192 | 0.060 |
| Residuals | 3275.034 | 28 | 116.966 | | | |
| stimulus | 1463.029 | 1 | 1463.029 | 0.332 | 0.569 | 0.012 |
| Residuals | 123476.404 | 28 | 4409.872 | | | |
| half | 757.812 | 1 | 757.812 | 0.199 | 0.659 | 0.007 |
| Residuals | 106703.372 | 28 | 3810.835 | | | |
| congruency ∗ stimulus | 66.597 | 1 | 66.597 | 0.360 | 0.553 | 0.013 |
| Residuals | 5173.092 | 28 | 184.753 | | | |
| congruency ∗ half | 622.414 | 1 | 622.414 | 2.583 | 0.119 | 0.084 |
| Residuals | 6747.709 | 28 | 240.990 | | | |
| stimulus ∗ half | 578.028 | 1 | 578.028 | 0.162 | 0.690 | 0.006 |
| Residuals | 99800.779 | 28 | 3564.314 | | | |
| congruency ∗ stimulus ∗ half | 24.440 | 1 | 24.440 | 0.112 | 0.741 | 0.004 |
| Residuals | 6118.941 | 28 | 218.534 | | | |
| Between subjects residuals | 597239.943 | 28 | 21329.998 | | | |

**Table S2.** Three way repeated measures ANOVA on RT data collected during the unmasked dot-probe task. The 2x2x2 design involves: *congruency* (the target presentation side congruent vs incongruent with respect to the self-/familiar-face presentation side), *stimulus* (self-face vs familiar-face), and *half* (first vs second half of trials within a block).

| source | Sum of Squares | df | Mean Square | F | p | $\eta_p^2$ |
|---|---|---|---|---|---|---|
| congruency | 825.593 | 1 | 825.593 | 3.781 | 0.062 | 0.119 |
| Residuals | 6114.681 | 28 | 218.381 | | | |
| stimulus | 61.811 | 1 | 61.811 | 0.037 | 0.850 | 0.001 |
| Residuals | 47393.982 | 28 | 1692.642 | | | |
| half | 838.851 | 1 | 838.851 | 0.849 | 0.365 | 0.029 |
| Residuals | 27672.339 | 28 | 988.298 | | | |
| congruency ∗ stimulus | 77.917 | 1 | 77.917 | 0.826 | 0.371 | 0.029 |
| Residuals | 2640.586 | 28 | 94.307 | | | |
| congruency ∗ half | 23.635 | 1 | 23.635 | 0.225 | 0.639 | 0.008 |
| Residuals | 2946.286 | 28 | 105.225 | | | |
| stimulus ∗ half | 725.099 | 1 | 725.099 | 1.041 | 0.316 | 0.036 |
| Residuals | 19495.681 | 28 | 696.274 | | | |
| congruency ∗ stimulus ∗ half | 1.097 | 1 | 1.097 | 0.009 | 0.923 | 0.000 |
| Residuals | 3247.771 | 28 | 115.992 | | | |
| Between subjects residuals | 337597.996 | 28 | 12057.071 | | | |

**Table S3.** Four way repeated measures ANOVA on ERP data collected during the masked dot-probe task. The 2x2x2x2 design involves: *N2pc* (signals recorded ipsi- vs contra-laterally with respect to the self-/familiar-face), *time-window* (early [200-300 ms] vs late [300-400 ms]), *stimulus* (self-face vs familiar-face), and *half* (first vs second half of experimental procedure).

| source | Sum of Squares | df | Mean Square | F | p | $\eta_p^2$ |
|---|---|---|---|---|---|---|
| N2pc | 1.490 | 1 | 1.490 | 4.931 | 0.035 | 0.150 |
| Residuals | 8.459 | 28 | 0.302 | | | |
| time-window | 108.722 | 1 | 108.722 | 21.540 | < .001 | 0.435 |
| Residuals | 141.329 | 28 | 5.047 | | | |
| stimulus | 0.012 | 1 | 0.012 | 0.002 | 0.961 | < .001 |
| Residuals | 142.883 | 28 | 5.103 | | | |
| half | 1.876 | 1 | 1.876 | 0.651 | 0.427 | 0.023 |
| Residuals | 80.700 | 28 | 2.882 | | | |
| N2pc ∗ time-window | 0.572 | 1 | 0.572 | 3.813 | 0.061 | 0.120 |
| Residuals | 4.199 | 28 | 0.150 | | | |
| N2pc ∗ stimulus | 0.116 | 1 | 0.116 | 0.344 | 0.562 | 0.012 |
| Residuals | 9.396 | 28 | 0.336 | | | |
| N2pc ∗ half | 1.302 | 1 | 1.302 | 1.744 | 0.197 | 0.059 |
| Residuals | 20.899 | 28 | 0.746 | | | |
| time-window ∗ stimulus | 3.963 | 1 | 3.963 | 6.621 | 0.016 | 0.191 |
| Residuals | 16.760 | 28 | 0.599 | | | |
| time-window ∗ half | 0.143 | 1 | 0.143 | 0.381 | 0.542 | 0.013 |
| Residuals | 10.527 | 28 | 0.376 | | | |
| stimulus ∗ half | 0.072 | 1 | 0.072 | 0.037 | 0.849 | 0.001 |
| Residuals | 54.436 | 28 | 1.944 | | | |
| N2pc ∗ time-window ∗ stimulus | 0.686 | 1 | 0.686 | 7.064 | 0.013 | 0.201 |
| Residuals | 2.719 | 28 | 0.097 | | | |
| N2pc ∗ time-window ∗ half | 0.199 | 1 | 0.199 | 2.685 | 0.112 | 0.088 |
| Residuals | 2.072 | 28 | 0.074 | | | |
| N2pc ∗ stimulus ∗ half | 1.528 | 1 | 1.528 | 3.634 | 0.067 | 0.115 |
| Residuals | 11.773 | 28 | 0.420 | | | |
| time-window ∗ stimulus ∗ half | 0.208 | 1 | 0.208 | 0.551 | 0.464 | 0.019 |
| Residuals | 10.566 | 28 | 0.377 | | | |
| N2pc ∗ time-window ∗ stimulus ∗ half | 0.024 | 1 | 0.024 | 0.346 | 0.561 | 0.012 |
| Residuals | 1.968 | 28 | 0.070 | | | |
| Between subjects residuals | 2564.108 | 28 | 91.575 | | | |

**Table S4.** Simple main effect of *N2pc* with a moderating factor of the *stimulus* and *time-window* for ERP data in the masked condition.

| Level of stimulus | Level of time-window | Sum of Squares | df | Mean Square | F | p | $\eta_p^2$ |
|---|---|---|---|---|---|---|---|
| self-face | early | 1.519 | 1 | 1.519 | 7.061 | 0.013 | 0.201 |
| Residuals | | 6.024 | 28 | 0.215 | | | |
| self-face | late | 0.124 | 1 | 0.124 | 0.657 | 0.424 | 0.023 |
| Residuals | | 5.278 | 28 | 0.189 | | | |
| familiar-face | early | 0.554 | 1 | 0.554 | 2.270 | 0.143 | 0.075 |
| Residuals | | 6.832 | 28 | 0.244 | | | |
| familiar-face | late | 0.666 | 1 | 0.666 | 2.810 | 0.105 | 0.091 |
| Residuals | | 6.639 | 28 | 0.237 | | | |

**Table S5.** Four way repeated measures ANOVA on ERP data collected during the unmasked dot-probe task. The 2x2x2x2 design involves: *N2pc* (signals recorded ipsi- vs contra-laterally with respect to the self-/familiar-face), *time-window* (early [200-300 ms] vs late [300-400 ms]), *stimulus* (self-face vs familiar-face), and *half* (first vs second half of experimental procedure).

| source | Sum of Squares | df | Mean Square | F | p | $\eta_p^2$ |
|---|---|---|---|---|---|---|
| N2pc | 7.216 | 1 | 7.216 | 7.312 | 0.012 | 0.207 |
| Residuals | 27.634 | 28 | 0.987 | | | |
| time-window | 257.972 | 1 | 257.972 | 39.248 | < .001 | 0.584 |
| Residuals | 184.040 | 28 | 6.573 | | | |
| stimulus | 0.030 | 1 | 0.030 | 0.005 | 0.947 | < .001 |
| Residuals | 187.593 | 28 | 6.700 | | | |
| half | 1.122 | 1 | 1.122 | 0.382 | 0.541 | 0.013 |
| Residuals | 82.155 | 28 | 2.934 | | | |
| N2pc ∗ time-window | 0.152 | 1 | 0.152 | 0.412 | 0.526 | 0.015 |
| Residuals | 10.346 | 28 | 0.369 | | | |
| N2pc ∗ stimulus | 11.646 | 1 | 11.646 | 14.151 | < .001 | 0.336 |
| Residuals | 23.042 | 28 | 0.823 | | | |
| N2pc ∗ half | 0.016 | 1 | 0.016 | 0.028 | 0.867 | 0.001 |
| Residuals | 15.321 | 28 | 0.547 | | | |
| time-window ∗ stimulus | 4.228 | 1 | 4.228 | 4.625 | 0.040 | 0.142 |
| Residuals | 25.596 | 28 | 0.914 | | | |
| time-window ∗ half | 0.570 | 1 | 0.570 | 1.145 | 0.294 | 0.039 |
| Residuals | 13.931 | 28 | 0.498 | | | |
| stimulus ∗ half | 0.334 | 1 | 0.334 | 0.156 | 0.696 | 0.006 |
| Residuals | 59.886 | 28 | 2.139 | | | |
| N2pc ∗ time-window ∗ stimulus | 0.411 | 1 | 0.411 | 1.598 | 0.217 | 0.054 |
| Residuals | 7.204 | 28 | 0.257 | | | |
| N2pc ∗ time-window ∗ half | 0.233 | 1 | 0.233 | 2.316 | 0.139 | 0.076 |
| Residuals | 2.812 | 28 | 0.100 | | | |
| N2pc ∗ stimulus ∗ half | < .001 | 1 | < .001 | < .001 | 0.987 | < .001 |
| Residuals | 20.989 | 28 | 0.750 | | | |
| time-window ∗ stimulus ∗ half | 2.439 | 1 | 2.439 | 5.403 | 0.028 | 0.162 |
| Residuals | 12.641 | 28 | 0.451 | | | |
| N2pc ∗ time-window ∗ stimulus ∗ half | 0.123 | 1 | 0.123 | 2.023 | 0.166 | 0.067 |
| Residuals | 1.704 | 28 | 0.061 | | | |
| Between subjects residuals | 2457.677 | 28 | 87.774 | | | |

**Table S6.** Simple main effect of *N2pc* with a moderating factor of the *stimulus* for ERP data in the unmasked condition.

| Level of stimulus | Sum of Squares | df | Mean Square | F | p | $\eta_p^2$ |
|---|---|---|---|---|---|---|
| self-face | 18.598 | 1 | 18.598 | 24.900 | < .001 | 0.471 |
| Residuals | 20.913 | 28 | 0.747 | | | |
| familiar-face | 0.264 | 1 | 0.264 | 0.248 | 0.622 | 0.009 |
| Residuals | 29.763 | 28 | 10.63 | | | |
